# Living cell-only bioink and photocurable supporting medium for printing and generation of engineered tissues with complex geometries

**DOI:** 10.1101/611525

**Authors:** Oju Jeon, Yu Bin Lee, Hyeon Jeong, Sang Jin Lee, Eben Alsberg

## Abstract

Scaffold-free engineering of three-dimensional (3D) tissue has focused on building sophisticated structures to achieve functional constructs. Although the development of advanced manufacturing techniques such as 3D printing has brought remarkable capabilities to the field of tissue engineering, technology to create and culture individual cell only-based high-resolution tissues, without an intervening biomaterial scaffold to maintain construct shape and architecture, has been unachievable to date. In this report, we introduce a cell printing platform which addresses the aforementioned challenge and permits 3D printing and long-term culture of a living cell-only bioink lacking a biomaterial carrier for functional tissue formation. A biodegradable and photocrosslinkable microgel supporting bath serves initially as a fluid, allowing free movement of the printing nozzle for high-resolution cell extrusion, while also presenting solid-like properties to sustain the structure of the printed constructs. The printed human stem cells, which are the only component of the bioink, couple together via transmembrane adhesion proteins and differentiate down tissue-specific lineages while being cultured in a further photocrosslinked supporting bath to form bone and cartilage tissue with precisely controlled structure. Collectively, this system, which is applicable to general 3D printing strategies, is paradigm shifting for printing of scaffold-free individual cells, cellular condensations and organoids, and may have far reaching impact in the fields of regenerative medicine, drug screening, and developmental biology.

## Introduction

Over the past decades, scaffolding approaches have been widely used to create functional tissues or organs in tissue engineering and regenerative medicine fields ^1^. However, the use of biomaterial-based scaffolds faces several challenges, such as interference with cell-cell interactions, potential immunogenicity of the materials and their degradation byproducts, unsynchronized rates of scaffold degradation with that of new tissue formation, and inhomogeneity and low density of seeded cells ^2^. To overcome these limitations of scaffold-based approaches, scaffold-free tissue engineering has recently emerged as a powerful strategy for constructing tissues using multicellular building blocks that self-assemble into geometries such as aggregates, sheets, strands and rings ^3^. These building blocks have been organized and fused into larger and more complicated structures, sometimes comprised of multiple cell types, and then they produce extracellular matrix (ECM) to form mechanically functional three-dimensional (3D) tissue constructs ^4-6^. However, it is still difficult to precisely control the architecture and organization of cell-only condensations to mimic sophisticated 3D structures of natural tissues and their structure-derived functions.

Recently, 3D printing has been applied in tissue engineering with the potential to create complicated 3D structures with high resolution using cell-free or cell-laden bioinks ^7^. Digital imaging data, obtained from computed tomography scans and magnetic resonance imaging, provide instruction for the desired geometry of printed constructs ^7,8^. Biodegradable thermoplastics, such as polycaprolactone, polylactic acid, and poly(lactic-co-glycolic acid), are advantageous for printing as stable constructs with delicate structural control can be formed due to the mechanical integrity of original materials ^9-11^. However, a major drawback is that cells cannot be printed simultaneously due to the use of organic solvents or high temperature to extrude the polymer inks ^12^. In contrast, materials that form cytocompatible and biocompatible hydrogels, such as alginate, gelatin, collagen, hyaluronic acid and polyethylene glycol, have been explored as prospective bioinks due to the feasibility of encapsulating cells within them during the printing process to provide a 3D cellular milieu ^7,13-15^. However, the hydrogel-based bioinks present the aforementioned limitations of scaffolding-based strategies. Since there has not been a platform enabling the printing and long-term culture of individual single cells while maintaining their resulting printed spatial position without incorporation of biomaterials, 3D cell printing in a scaffold-free manner has rarely been achieved. Bhattacharjee et al. reported living-cell only bioinks. However, their supporting medium could not provide long-term support for 3D printed structures ^16^. Instead of single cell-based bioprinting, there is a platform that permits skewering pre-cultured and formed multicellular aggregates on an array of needles to assemble structures of interest ^17^. Cell strands, which are acquired by pre-culture and assembly of cells in a hollow fiber mold, have also been used as bioink for scaffold-free 3D bioprinting ^18^. For these examples, the resolutions of the systems are limited by the original size and shape of the bioink cellular assemblies, preparing the bioinks demands additional time and expense, and neither permits printing of individual cells. Moreover, complicated instrumental setups required, such as vacuum syringe assisted skewering, may limit widespread implementation due to associated costs and necessary expertise.

Here, we present newly generated tissues from directly assembled stem cells, which have been 3D bioprinted into a photo-curable liquid-like supporting medium comprised of solid hydrogel microparticles (microgels) (Fig. S1). The supporting bath consists of biodegradable and photocrosslinkable alginate microgels, which are prepared by ionic crosslinking of dual-crosslinkable, oxidized and methacrylated alginate (OMA) ^19^, and is expected to applicable to general 3D bioprinting systems. The microgel supporting medium sustains the high-resolution printing of human bone marrow-derived mesenchymal stem cells (hMSCs) by exhibiting similar properties to Bingham plastic fluids ^20^. While the microgel supporting medium allows the printing needle move freely via its shear-thinning properties, the microgels work as supporting materials for printed constructs through self-healing properties ^21^. After directly 3D bioprinting of hMSCs into the microgel supporting medium, photocrosslinking of the microgels can provide mechanical stability for hMSC constructs for long-term culture. Dissociation of the photocrosslinked microgel supporting medium by gentle agitation may facilitate acquisition of matured 3D tissue constructs. Collectively, our objectives were (i) to assess the effect of the size of dual-crosslinkable OMA microgels in the supporting bath on printing resolution, (ii) to evaluate the capacity of the OMA supporting bath to maintain the viability individual printed cells and the structure of resulting self-assembly printed constructs, and (iii) to investigate function of the obtained 3D scaffold-free cellular constructs.

## Results

### 3D bioprinting of living hMSCs without a biomaterial in the bioink

Living hMSCs can be printed as a bioink by themselves without a carrier macromer solution into a photo-curable, self-healing and shear-thinning alginate microgel supporting medium, which is formed with calcium-crosslinked OMA microgels (Fig. 1). Alginate microgel supporting medium is fluidized under low shear stress, permitting easy insertion and rapid motion of needles deep within the bulk. After removing shear stress caused by needle movement and ejection of printing material, the locally fluidized alginate microgel bath rapidly “self-heals” and forms a stable medium that firmly holds the printed hMSCs in 3D place (Fig. 1a). The low yield stress of the alginate microgel medium in its solid state and its rapid self-healing behavior allows the unrestricted deposition, placement and structuring of cells deep within the alginate microgel supporting medium that maintains the printed structure with fidelity (Fig. 1b and Movie S1 and S2; The movies play at 4x speed.). To explore the versatility and stability of 3D printing into the alginate microgel supporting medium, a variety of complicated 3D structures were printed using only individual cells as a bioink. A letter (C), an ear, letters comprising an acronym (CWRU) and a femur were successfully created with high resolution (Fig. 1c-f).

**Fig. 1.**
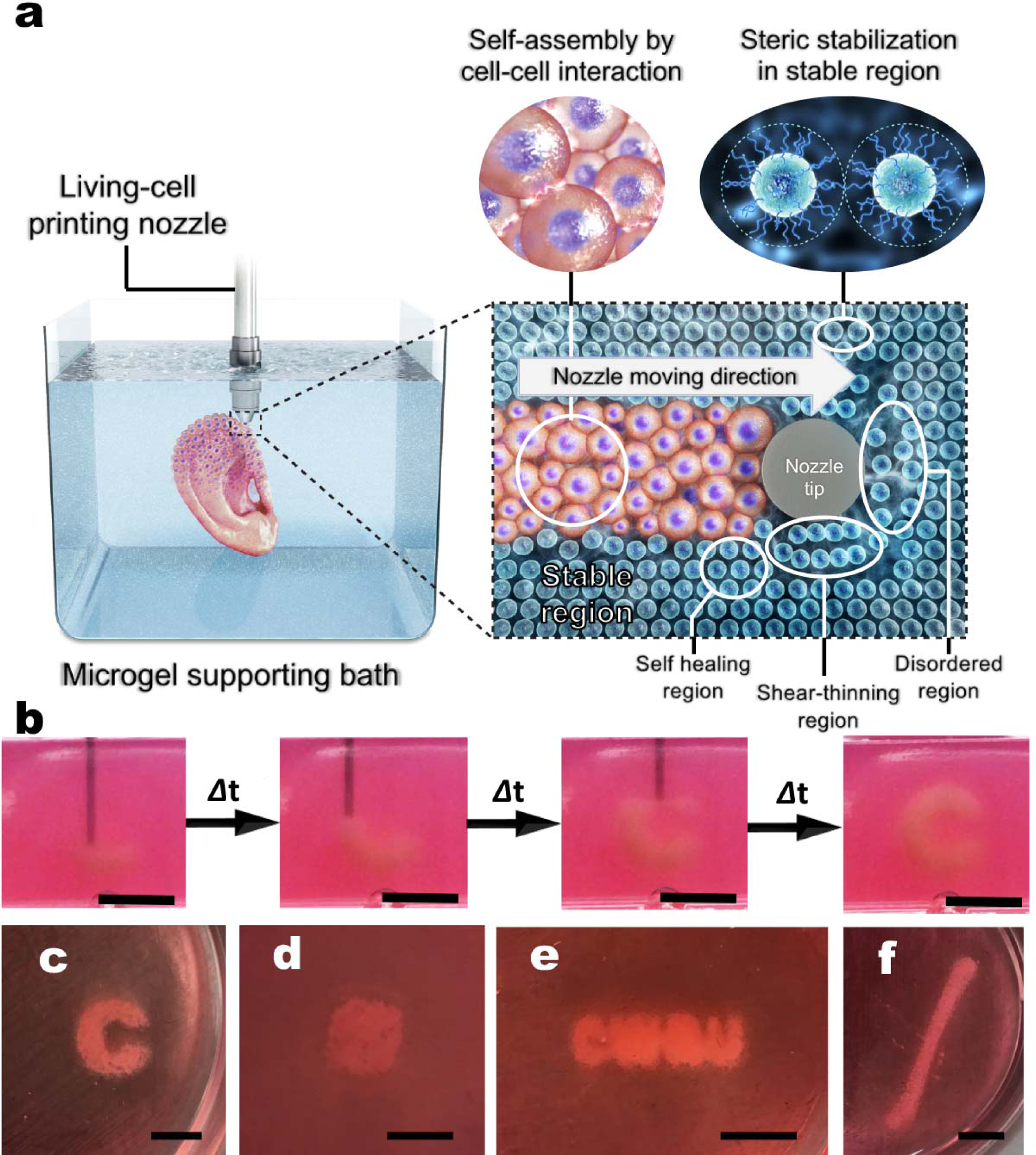
Shear-thinning and self-healing alginate microgel supporting medium for 3D bioprinting of living stem cells. **(a)** A schematic of 3D printing of cells within the alginate microgel supporting medium. OMA microgels in the supporting medium fluidize via their shear-thinning properties when stress is applied by motion of the printing needle and cell-only bioink (shear-thinning region) and rapidly fill in after the needle passes by self-healing properties (self-healing region) without creating crevasses. Microgel supporting medium without shear stress presents solid-like properties, which provide mechanical stability for the printed cell construct (stable region). **(b)** Captured images at different times during bioprinting of the letter “C” using living stem cell-only bioink into the alginate microgel supporting medium. As the printing progress, cells are arranged into the letter “C” shape in 3D without disturbing previously printed regions, which is achieved as a result of the shear-thinning and self-healing properties of the alginate microgel supporting medium. **(c)** Images of the 3D bioprinted structures of a letter “C”, a cube, letters comprising the acronym “CWRU”, and a femur in alginate microgel supporting medium. Scale bars indicate 5 mm.

### Properties of the alginate microgel supporting medium

To identify favorable properties of alginate microgels for use as supporting medium for 3D cell printing, several rheological measurements were performed on supporting medium made up of two different sizes of alginate microgels (Fig. 2 and Fig. S2). To verify the solid-like properties of alginate microgel supporting medium, a frequency sweep at low strain amplitude (1%) was conducted, measuring the elastic and viscous shear moduli and viscosity. The data show both sizes (7.0 ± 2.8 and 409.5 ± 193.7*μ*m, Fig. S1) of alginate microgels behave like solid materials at low shear strain due to the steric stabilization of highly packed microgels (Fig. 2a and Fig. S2a) ^22^, but they exhibit shear-thinning properties with decreased viscosity as shear rate increases (Fig. 2b and Fig S2b). To further identify the shear-thinning and shear yielding properties of the alginate microgel supporting medium in response to shear strain, the shear moduli with a strain sweep at a constant frequency (1Hz) was measured. Both sizes of OMA microgels exhibited shear-thinning (Fig. 2c and Fig. S2c) and shear-yielding (Fig. 2d and Fig. S2d) properties following increased shear strain application. Although both sizes of microgels exhibited a crossover at similar strain amplitude, the modulus at the crossover point (G’ = G”) of the smaller OMA microgels was much lower than that of the larger OMA microgels (Fig. 2d and Fig. S2d). To characterize the self-healing or recovery behavior of the alginate microgel medium, dynamic strain tests were performed with alternate low (1%) and high (100%) strains. A rapid recovery of the storage modulus (Fig. 2e and Fig. S2e) and viscosity (Fig. 2f and Fig. S2f) within seconds to the initial properties was repeatedly achieved over several cycles for both sizes of alginate microgels, indicating that the alginate microgel supporting medium can rapidly change from the solid to the fluid state via application of shear strain. Printing materials into viscoelastic supporting materials often results in crevasses created by the movement of the shaft of the dispensing needle and requires a third material that fills in crevasses.^23^ However, 3D structures of hMSCs can be written into alginate microgel supporting medium without creating crevasses due to the self-healing properties of the alginate microgel supporting medium (Movie S3). To evaluate the capacity of the OMA microgels to provide long-term support for 3D printed constructs, frequency (at 1 % strain) and strain (at 1Hz) sweep tests were conducted after photocrosslinking of the smaller sized OMA microgel-based supporting medium under low-level UV light. Frequency (Fig. 2g) and strain (Fig. 2h) sweeps exhibited significantly higher G’ than G’’, indicating that photocrosslinked OMA microgel supporting medium is mechanically stable without shear yielding. The stability of photocrosslinked OMA microgel supporting medium was also confirmed by a wash out test (Fig. S3). While the photocrosslinked OMA microgel supporting medium remained stable on the Petri dish, uncrosslinked OMA microgel supporting medium could be easily removed by washing with water.

**Fig. 2.**
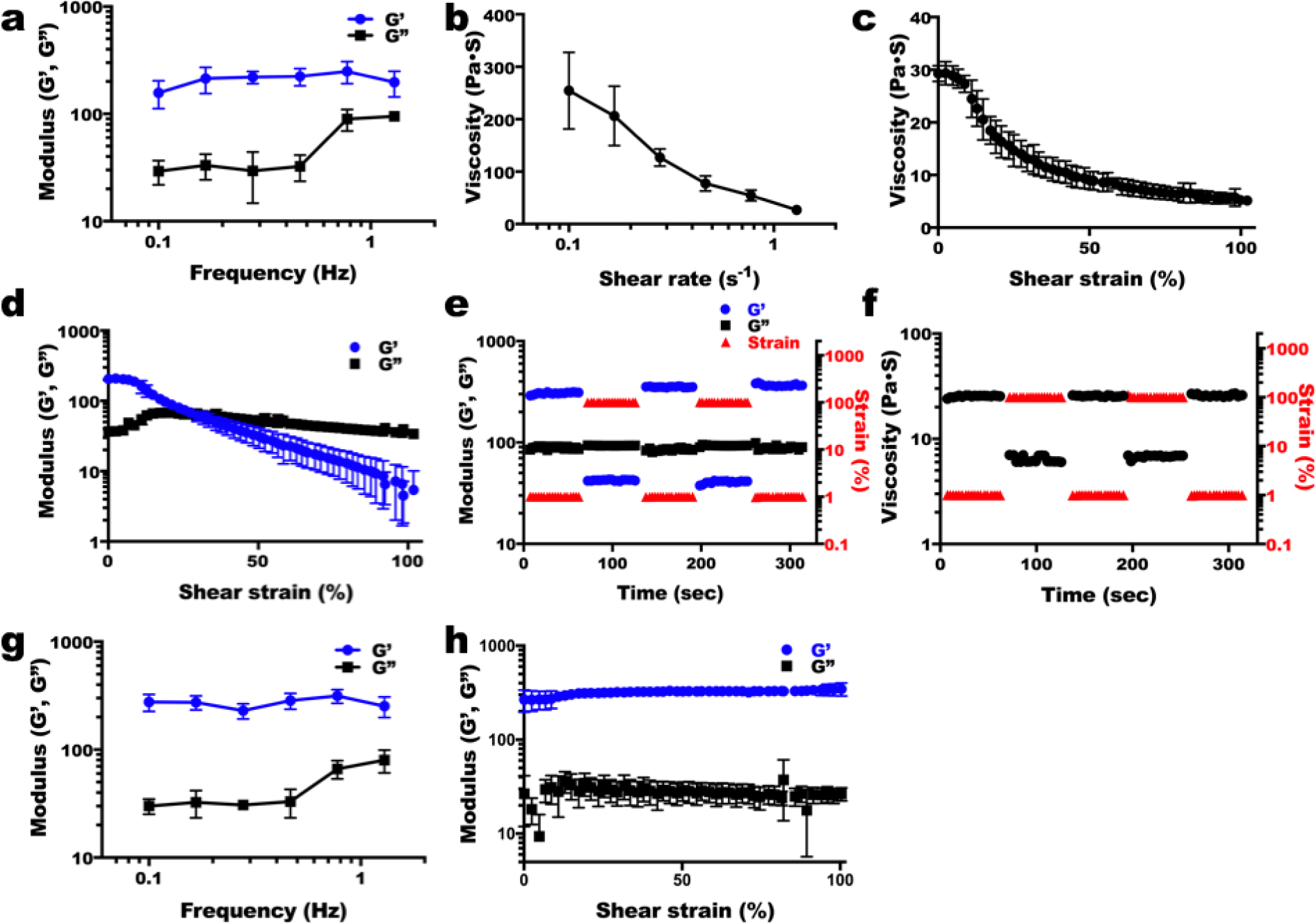
Shear-thinning and self-healing properties of the alginate microgel supporting medium. **(a)** Storage (G’) and loss (G”) moduli of alginate microgel supporting medium (mean microgel diameter = 7.0 ± 2.8 μm) as a function of frequency. G’ is larger than G” over the measured frequency range and both moduli exhibit frequency independence. Viscosity measurements of alginate microgel supporting medium as a function of **(b)** shear rate and **(c)** shear strain demonstrate its shear-thinning behavior. **(d)** G’ and G” of the alginate microgel supporting medium as a function of shear strain exhibit its shear-yielding behavior and gel-to-sol transition at higher shear strain. **(e)** Shear moduli and **(f)** viscosity changes in dynamic strain tests of the alginate microgel supporting medium with alternating low (1%) and high (100%) strains at 1 Hz demonstrate its rapid recovery of strength and viscosity within seconds, which indicates “self-healing” or thixotropic properties. **(g)** Frequency sweep (at 1 % strain) and **(h)** strain sweep (at 1 Hz) tests of the alginate microgel supporting medium after photocrosslinking under low-level UV light. G’ is larger than G’’ over the measured frequency and strain ranges and both moduli exhibit frequency and strain independence, indicating that the photocrosslinked alginate microgel supporting medium is mechanically stable.

### Characterization of 3D printed cell-only filaments

Next, it was important to determine the minimum printed structure feature size achievable using this strategy. Lines or “filaments” of cells were printed into supporting medium with both sizes of alginate microgels to compare resulting resolutions. Regardless of the microgel size, hMSCs in filaments exhibited high cell viability as visualized by live/dead assay, demonstrating no adverse effects of the bioprinting process and UV irradiation for curing the microgel supporting medium on cell survival (Fig. 3a-c and e-f). The smaller alginate microgel supporting medium (Fig. 3d) exhibited higher resolution with narrow filament diameter distribution compared to the larger alginate microgel supporting medium (Fig. 3h), while the mean diameters of both hMSC filaments were similar (395.1 ± 64.6 and 419.8 ± 187.5 *μ*m for filaments printed in small and larger microgel supporting medium, respectively). Since medium pores result from the space between the microgels, larger microgels make larger medium pores and vice versa. Due to the larger pores, many hMSCs printed into the larger alginate microgel supporting medium dispersed into the medium from the filaments, while hMSC filaments printed into the smaller alginate microgel supporting medium show a limited dispersion of cells. Therefore, 3D printed hMSC constructs in the smaller alginate microgel supporting medium (Fig. 3i and j) exhibit higher resolution than those in the larger alginate microgel supporting medium (Fig. 3k and l). These results indicate that the supporting medium comprised of smaller alginate microgels, which has lower stiffness, yield strength and viscosity, is more favorable for printing hMSCs with high resolution. Importantly, when cells were printed into the smaller alginate microgel supporting medium with smaller-gauge needles (25 and 27 G), significantly higher resolution of hMSC filaments (p<0.05, one-way ANOVA with Tukey’s multiple comparison test using GraphPad Prism) was achieved (Fig. 3m-r) compared to that with the larger-gauge needle (Fig. 3a-c).

**Fig. 3.**
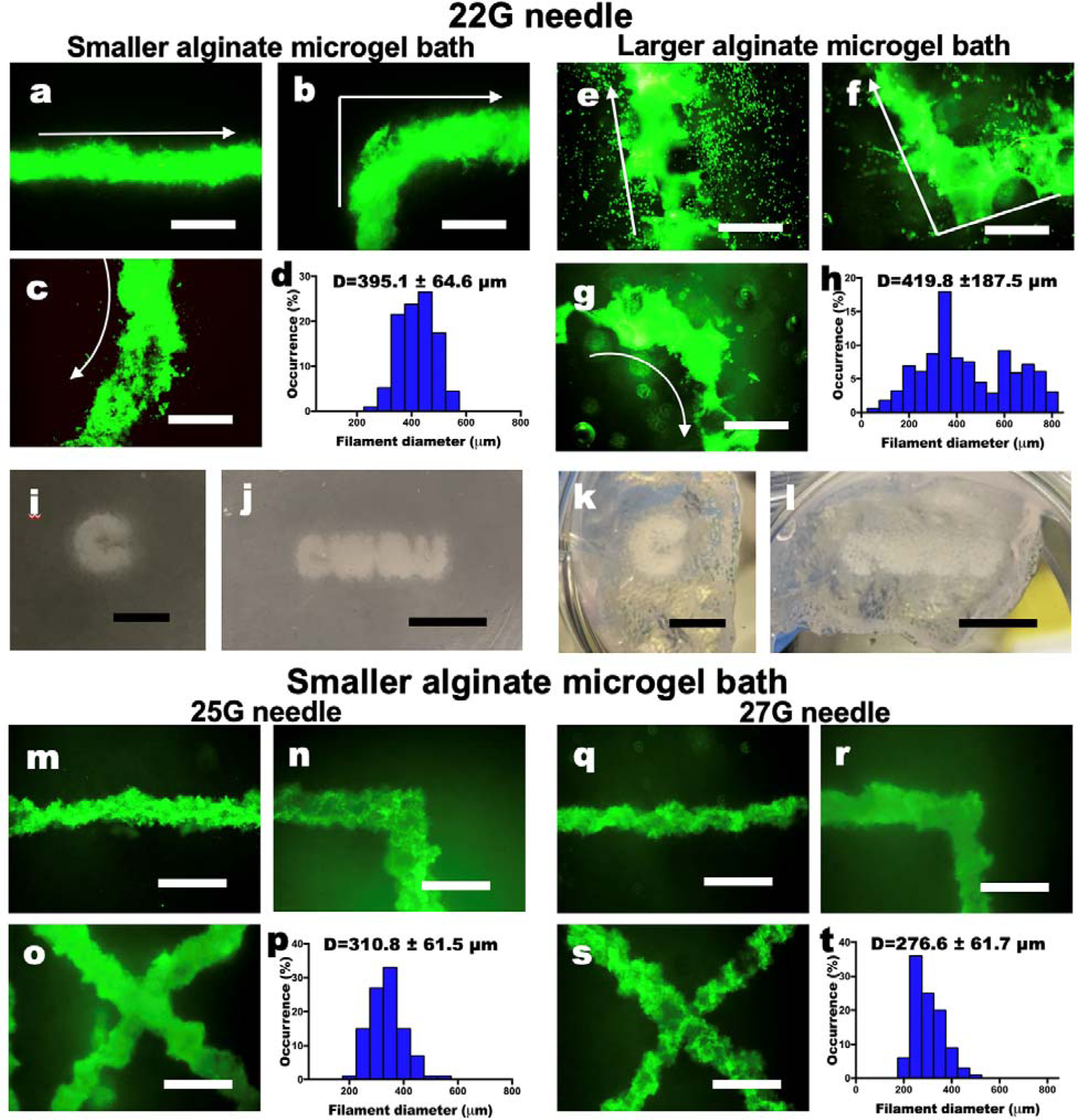
Characterization of living cell bioink. **(a-c)** Live/Dead staining of 3D hMSC filaments bioprinted in a straight line, a corner and a curve with a **22 G** needle and **(d)** their diameter distribution in **the smaller alginate microgel supporting medium**. **(e-g)** Live/Dead staining of 3D hMSC filaments bioprinted in various configurations with a **22 G** needle and **(h)** their diameter distribution in **the larger alginate microgel supporting medium**. Arrows indicate the direction of movement of the printing nozzle. Scale bars indicate 600 *μ*m. The Live/Dead images demonstrate high cell viability. Smaller alginate microgels lead to higher resolution printing by limiting diffusion of cells into the pores of the microgel bath. Thickness of the cell filaments also are more narrowly distributed in smaller microgel medium. Images of letters ‘C’ and “CWRU” in **(i and j)** the smaller and (**k and l)** larger alginate microgel supporting medium after photocrosslinking. Scale bars indicate 5 mm. **(m-o)** Live/Dead staining of 3D hMSC filaments bioprinted in various configurations with a **25 G** needle and **(p)** their diameter distribution in **the smaller alginate microgel supporting medium**. **(q-s)** Live/Dead staining of 3D hMSC filaments bioprinted in various configurations with a **27 G** needle and **(h)** their diameter distribution in **the smaller alginate microgel supporting medium**. Scale bars indicate 600 *μ*m. Smaller diameter needles lead to higher resolution printing of the cell filaments, which also are more narrowly distributed. Scale bars indicate 600 *μ*m.

### 3D printing of complex structures and formation of engineered tissues

Long-term cell culture is essential to ensure tissue formation through maintenance cell-cell interactions, self-assembly into cellular condensations, and differentiation of the stem cells down desired lineages for engineering specific tissue types ^24^. Critical to achieving this for a cell-only bioink is the capacity to provide the mechanical stability with the supporting medium during the culture period. Since the alginate microgels possess photo-reactive methacrylate groups, the medium can be further photocrosslinked to form a more stable supporting structure that retains its shape for extended culture. After photocrosslinking, the alginate microgel supporting medium exhibited robust mechanical stability without shear yielding (Fig. 2h), maintained initial 3D printed structures (Fig. 1c-f) and enabled long-term culture of 3D printed constructs for formation of functional tissue by differentiation of 3D bioprinted hMSCs. After 4 weeks of osteogenic or chondrogenic differentiation, formed tissue constructs were easily harvested from the alginate microgel supporting medium by applying shear force using a pipette. 3D printed hMSCs were assembled into precise multicellular structures following the architecture defined by computer-aided design (CAD) files (Fig. 4a and d), and bone- (Fig. 4b-c) or cartilage- (Fig. 4e-f and Fig. S4) like tissues were obtained in the photocrosslinked alginate microgel supporting medium. Differentiation down the osteogenic and chondrogenic lineages and resultant formation of bone and cartilage tissue were confirmed via Alizarin red (red) and Toluidine blue O (purple) staining, respectively; red and purple colors were intensively observed throughout the constructs (Fig. 4c and f) and sectioned samples (Fig. 4g and h). Lacunae structures were also observed in sectioned slides of chondrogenically differentiated constructs (Fig. 4h), indicating maturation of cartilage tissues ^25^. Successful tissue formation by the 3D printed hMSCs were further confirmed by quantification of osteogenic (i.e., alkaline phosphatase (ALP) activity and calcium deposition) and chondrogenic (i.e., glycosaminoglycan (GAG) production) markers (Fig. S5). Collectively, the microgel supporting medium allows not only high-resolution printing of cell-only bioink, but also provides printed construct mechanical stability after additional photocrosslinking, which permits culture of the constructs with stable structural maintenance and long-term differentiation in differentiation medium.

**Fig. 4.**
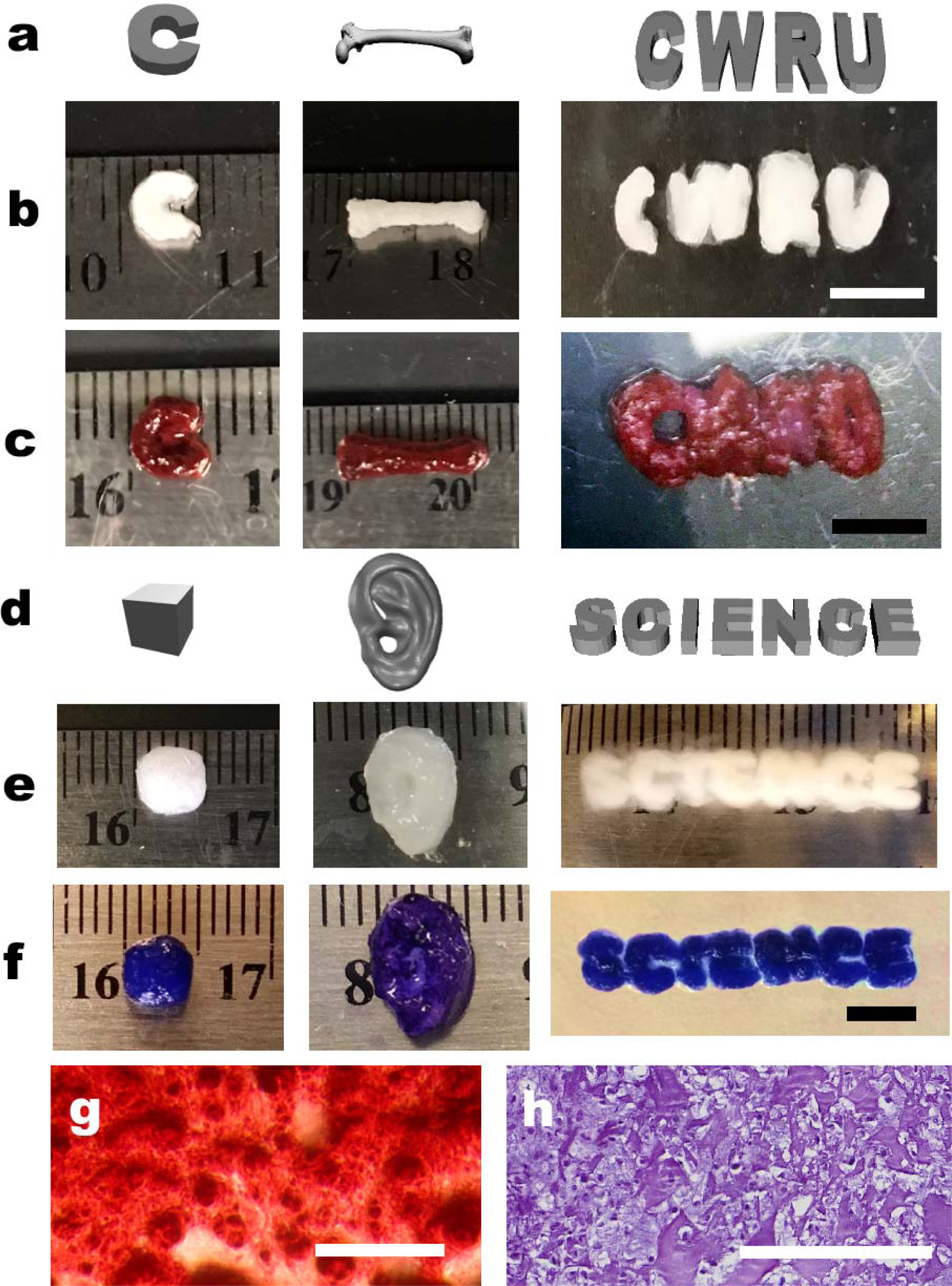
Differentiation of 3D bioprinted hMSC constructs. **(a)** Digital images and photographs of osteogenically differentiated 3D printed hMSC construct morphology **(b)** before and **(c)** after Alizarin red S staining. Scale bars indicate 5 mm. **(d)** Digital images and photographs of chondrogenically differentiated 3D printed hMSC construct morphology **e)** before and **(f)** after Toluidine blue O staining. The constructs presented well-preserved structures after long-term 4-week culture without evidence of construct deformation due to cellular contraction or proliferation, and generation of specific tissue types (i.e., bone and cartilage) with desired geometries. Scale bar indicates 5 mm. Photomicrographs of **(g)** Alizarin Red S and **(h)** Toluidine Blue O stained construct sections. The images demonstrate hMSC differentiation and deposition of lineage specific ECM in the cell-only bioink printed constructs. Scale bars indicate 200 *μ*m.

## Discussion

As we describe in this study, 3D tissue structures can be created using single cell-based bioprinting with an assistance of OMA microgel supporting medium for precise structural control and long-term culture for formation of functional engineered tissues. Since the platform is applicable to universal 3D printers, it doesn’t demand experts in either software or hardware fields ^20^. Using a printing set-up costing <$1K, rapid construct formation on the centimeter scale was achieved in a few minutes, and even faster and more complex tissue printing with higher resolution may be accomplished with higher quality printers. As the microgel size decreased, it was possible to build tissue constructs with sophisticated structures due to the medium shear-thinning properties upon needle motion, self-healing properties in absence of external strain, and limited diffusion of the printed cells into its pores ^26^. In addition, high viability of the printed cells was realized even after additional photocrosslinking of the microgels for long-term structural support. Unlike previous 3D bioprinting techniques which depend on external solid materials for structural maintenance or additional process for prefabrication of cell aggregates ^17^, the photocrosslinked OMA microgel supporting medium played a structural support role for the printed cell constructs, allowing media provision and long-term culture. Precise maintenance of the structure, mirroring the original CAD file design, was also achieved even after maturation of the tissue which possibly caused deformation, shrinking and/or thickening of the printed constructs due to cell proliferation, differentiation and ECM production ^27^. Since the OMA microparticles can be removed by simple agitation or spontaneous degradation from the constructs, the cultured constructs can be easily harvested from the alginate microgel supporting medium without damage. This universally applicable 3D printing platform makes it possible to print isolated cells without a biomaterial carrier in the bioink, and will contribute to regenerative medicine by permitting generation of biomimetic cellular condensation-based engineered tissues with defined geometries comprised of multiple cell types with controlled spatial placement.

## Materials and Methods

Complete detailed methodology can be found in Supporting Information.

## Acknowledgments

The authors gratefully acknowledge funding from the National Institutes of Health’s National Institute of Arthritis And Musculoskeletal and Skin Diseases under award numbers R01AR063194 and R01AR066193 and National Institute of Biomedical Imaging and Bioengineering under award number R01EB023907. The contents of this publication are solely the responsibility of the authors and do not necessarily represent the official views of the National Institutes of Health.

## Data availability

The data that support the findings of this study are available from the corresponding author upon reasonable request.

## Supporting Information

### Materials and Methods

#### Synthesis of OMA

Oxidized alginate (OA) was prepared by reacting sodium alginate with sodium periodate^19^. Sodium alginate (10 g, Protanal LF 200S, FMC Biopolymer) was dissolved in ultrapure deionized water (diH_2_O, 900 ml) overnight. Sodium periodate (0.1 g, Sigma) was dissolved in 100 ml diH_2_O, added to alginate solution under stirring to achieve 1 % theoretical alginate oxidation, and allowed to react in the dark at room temperature for 24 hrs. The oxidized, methacrylated alginate (OMA) macromer was prepared by reacting OA with 2-aminoethyl methacrylate (AEMA). To synthesize OMA, 2-morpholinoethanesulfonic acid (MES, 19.52 g, Sigma) and NaCl (17.53 g) were directly added to an OA solution (1 L) and the pH was adjusted to 6.5. N-hydroxysuccinimide (NHS, 1.176 g, Sigma) and 1-ethyl-3-(3-dimethylaminopropyl)-carbodiimide hydrochloride (EDC, 3.888 g, Sigma) were added to the mixture under stirring to activate 20 % of the carboxylic acid groups of the alginate. After 5 min, AEMA (1.688 g, Polysciences) (molar ratio of NHS:EDC:AEMA = 1:2:1) was added to the solution, and the reaction was maintained in the dark at RT for 24 hrs. The reaction mixture was precipitated into excess of acetone, dried in a fume hood, and rehydrated to a 1 % w/v solution in diH_2_O for further purification. The OMA was purified by dialysis against diH_2_O using a dialysis membrane (MWCO 3500, Spectrum Laboratories Inc.) for 3 days, treated with activated charcoal (5 g/L, 50-200 mesh, Fisher) for 30 min, filtered (0.22 μm filter) and lyophilized. To determine the levels of alginate methacrylation, the OMA was dissolved in deuterium oxide (D_2_O, 2 w/v %), and ^1^H-NMR spectra were recorded on a Varian Unity-300 (300MHz) NMR spectrometer (Varian Inc.) using 3-(trimethylsilyl)propionic acid-d_4_ sodium salt (0.05 w/v %) as an internal standard.

#### Fabrication of OMA microgel slurry

To fabricate smaller OMA microgels, OMA (1.5 g) was dissolved in DMEM (100 ml) containing 0.05 % photoinitiator (PI), placed in a syringe, and then dispensed into a gelling bath containing an aqueous solution of CaCl_2_ (1 L, 0.2 M) bath. After fully ionically crossliking the OMA fibers in the bath for 30 min, the resultant OMA fibers were collected, washed with DMEM three times, and then blended using a consumer-grade blender (Osterizer MFG, at “pulse” speed) for 90 sec with 100 ml DMEM. Then, the blended OMA slurry was loaded into 50 ml conical tubes and centrifuged at 2000×g for 5 min. The supernatant was removed and replaced with a sterile 70 % ethanol. The slurry was vortexed back into suspension and centrifuged again. After the supernatant was removed, the OMA microgel slurry was vortexed with sterile 70 % ethanol and then stored until use at 4 °C. To fabricate larger sized OMA micogels, OMA solution was loaded into a 3-ml syringe, and then the syringe was connected to a custom coaxial microdroplet generator designed in our laboratory (Fig. S6). The OMA solution was pumped at 0.5 ml/sec with an outer air flow rate of 10 L/min, and the droplets dripped into a collection bath containing an aqueous solution of CaCl_2_ (0.2 M). After fully ionically crosslinking the microgels in the bath for 30 min, the resultant OMA microgels were collected and washed with DMEM three times. The OMA microgels were suspened in sterile 70 % ethanol and stored until use at 4 °C.

To evaluate the morphology and measure the size of OMA microgels comprising the slurries, the slurries were centrifuged at 2000×g for 5 min. The supernatants were removed and replaced with DMEM containing 0.05 % PI, and the microgels were vortexed back into suspension and then centrifuged again. This process was repeated five times and then the supernatants were removed. To visualize the OMA microgels, they were stained with Safranin O and then imaged using a microscope (Leitz Laborlus S, Leica) equipped with a digital camera (Coolpix 995, Nikon). To measure the mean diameter of the smaller OMA microgels (prepared using a blender), 1 ml of the OMA microgels were suspended in 10 ml DMEM containing 0.05 % PI and measured at room temperature by dynamic light scattering using a particle size analyzer (90Plus, Brookhaven Instruments). The mean diameter of the larger OMA microgels (prepared via the coaxial microdroplet generator) was measured using ImageJ with the images of Safranin O stained OMA microgels.

#### Rheological properties of OMA microgel slurry

Dynamic rheological examination of the OMA microgel slurries was performed to evaluate shear-thinning, self-healing and mechanical properties with a Haake MARS III rotational rheometer (ThermoFisher Scientific). In oscillatory mode, a parallel plate (80 mm diameter) geometry measuring system was employed, and the gap was set to 1 mm. After each OMA microgel slurry was placed between the plates, all the tests were started at 37 ± 0.1 °C, and the plate tempearature was maintained at 37 °C. Oscillatory frequency sweep (0.01-1.3 Hz at 1 % strain) tests were performed to measure storage moduli (G’), loss moduli (G”) and viscosity. Oscillatory strain sweep (0.1-100 % strain at 1 Hz) tests were performed to show the shear-thinning characteristics of the OMA microgels and to determine the shear-yielding points at which the OMA microgel slurries behave fluid-like. To demonstrate the self-healing properties of OMA microgel slurries, cyclic deformation tests were performed at 100 % strain with recovery at 1 % strain, each for 1 min at 1 Hz.

#### Preparation of hMSC ink

To isolate human bone marrow-derived mesenchymal stem cells (hMSCs) ^28^, bone marrow aspirates were obtained from the posterior iliac crest of a healthy twenty seven-year old male donor under a protocol approved by the University Hospitals of Cleveland Institutional Review Board. The aspirates were washed with growth medium comprised of low-glucose Dulbecco’s Modified Eagle’s Medium (DMEM-LG, Sigma) with 10 % prescreened fetal bovine serum (FBS, Gibco). Mononuclear cells were isolated by centrifugation in a Percoll (Sigma) density gradient and the isolated cells were plated at 1.8 x 10^5^ cells/cm^2^ in DMEM-LG containing 10 % FBS and 1 % penicillin/streptomycin (P/S, Thermo Fisher Scientific) in an incubator at 37 °C and 5 % CO_2_. After 4 days of incubation, non-adherent cells were removed and adherent cell were maintained in DMEM-LG containing 10 % FBS, 1 % P/S and 10 ng/ml FGF-2 with media changes every 3 days. After 14 days of culture, the cells were passaged at a density of 5×10^3^ cells/cm^2^, cultured for an additional 14 days, and then stored in cryopreservation media in liquid nitrogen until use. To use hMSC as a bioink, hMSCs were expanded in growth media consisting of DMEM-LG with 10% FBS (Sigma), 1 % P/S and 10 ng/ml FGF-2 and loaded into a 2.5-ml syringe (Gastight Syringe, Hamilton Company).

#### Modification of 3D printer

All 3D printing was performed using a 3D printer (PrintrBot Simple Metal 3D Printer, Printrbot) modified with a syringe-based extruder ^14^. The stock thermoplastic extruder assembly was replaced with a custom-built syringe pump extruder. The syringe pump extruder was designed to use the NEMA-17 stepper motor from the original Printrbot thermoplastic extruder and mount directly in place of the extruder on the *x*-axis carriage. The syringe pump extruder was printed with polylactic acid using the thermoplastic extruder on the Printrbot before its removal. Using the same stepper motor, the syringe pump extruder was natively supported by the software that came with the printer. The design for the syringe pump extruder and the image file of the human femur were downloaded as STL files from the NIH 3D Print Exchange (http://3dprint.nih.gov) under open-source license. Digital image files of letters for 3D printing were generated from www.tinkercad.com. The file for the human ear was downloaded from www.thinkiver.com/thing:304657 under the terms of the Creative Commons Attribution license, which permits unrestricted use, reproduction and distribution in any medium.

#### 3D printing of hMSCs

The hMSC-loaded syringe was connected to a 0.5-inch 22G stainless steel needle (McMaster-Carr) and mounted into the syringe pump extruder on the 3D printer. A petri dish was filled with OMA microgel slurry at room temperature to serve as a supporting bath and placed on the building platform. The tip of the needle was positioned at the center and near the bottom of the dish, and the print instructions were sent to the printer using the host software (Cura Software, Ultimaker), which is an open source 3D printer host software. After 3D printing of hMSCs, OMA microgel supporting medium with a 3D printed construct was stabilized by photocrosslinking under UV at 20 mW/cm^2^ for 1 min. After slurry photocrosslinking, 3D printed hMSC constructs in the photocrosslinked OMA microgel slurry were transferred into 6-well tissue culture plates with growth media, chondrogenic differentiation media or osteogenic differentiation media, and placed in a humidified incubator at 37 °C with 5 % CO_2_.

#### Analysis of printed hMSC structures

Linear hMSC filaments were printed in the OMA microgel supporting baths with 22, 25 and 27 G needles, baths were photocrosslinked under UV light a 20 mW/cm^2^ for 1 min, and then 5 ml culture media was added. The viability and morphology of 3D printed hMSC filaments were investigated using a Live/Dead staining comprised of fluorescein diacetate [FDA, 1.5 mg/ml in dimethyl sulfoxide (Research Organic Inc.), Sigma] and ethidium bromide (EB, 1 mg/ml in PBS, Thermo Fisher Scientific). The staining solution was freshly prepared by mixing 1 ml FDA solution and 0.5 ml EB solution with 0.3 ml PBS (pH 8). 100 μl of staining solution was added into each well and incubated for 10 min at room temperature, and then stained 3D printed hMSC filaments were imaged using a fluorescence microscope (ECLIPSE TE 300) equipped with a digital camera (Retiga-SRV). Diameters of the 3D printed hMSC filaments were measured at least 400 times for each group using ImageJ (National Institutes of Health).

#### Osteogenic and chondrogenic differentiation of the 3D printed hMSC constructs

3D printed hMSC constructs in the photocrosslinked OMA microgel supporting baths were differentiated by culture with osteogenic differentiation media [10 mM β-glycerophosphate (CalBiochem), 37.5 μg/ml ascorbic acid (Wako USA), 100 nM dexamethasone (MP Biomedicals), and 100 ng/ml BMP-2 in DMEM-high glucose] containing 10 % FBS and 1% P/S or chondrogenic differentiation media [1 % ITS+ Premix, 100 nM dexamethasone, 37.5 μg/ml l-ascorbic acid-2-phosphate, 1 mM sodium pyruvate, 100 μM nonessential amino acids, and 10 ng/ml TGF-β_1_ in DMEM-high glucose]. The osteogenic and chondrogenic media was changed twice a week. After 4 weeks of culture in osteogenic differentiation media, 3D printed hMSC constructs were fixed in 10 % neutral buffered formalin overnight at 4 °C and stained with Alizarin red S. Cryosectioned samples were also stained with Alizarin red S. After 3 weeks of culture in chondrogenic differentiation media, 3D printed hMSC constructs were fixed in 10 % neutral buffered formalin over night at 4 °C and stained with Toluidine blue O. Cryosectioned samples were also stained with Toluidine blue O. For quantification of alkaline phosphatase (ALP) activity, DNA content and calcium deposition, osteogenically differentiated 3D printed hMSC constructs were homogenized at 35,000 rpm for 30 s using a TH homogenizer (Omni International) in 1ml ALP lysis buffer (CelLytic™ M, Sigma). The homogenized solutions were centrifuged at 500 g with a Sorvall Legent RT Plus Centrifuge (Thermo Fisher Scientific). For ALP activity measurements, supernatant (100 μl) was treated with p-nitrophenylphosphate ALP substrate (pNPP, 100 μl, Sigma) at 37 °C for 30 min, and then 0.1 N NaOH (50 μl) was added to stop the reaction. The absorbance was measured at 405 nm using a plate reader (FMAX, Molecular Devices) (N=4). A standard curve was made using the known concentrations of 4-nitrophenol (Sigma). DNA content in supernatant (100 μl) was measured using a Quant-iT PicoGreen assay kit (Invitrogen) according to the manufacturer’s instructions. Fluorescence intensity of the dye-conjugated DNA solution was measured using a plate reader (FMAX) set at 485 nm excitation and 538 nm emission (N=4). After an equal volume of 1.2 N HCl was added into each lysate solution, the mixed solutions were centrifuged at 500 g with a Sorvall Legent RT Plus Centrifuge. Calcium deposition of the constructs was quantified using a calcium assay kit (Pointe Scientific) according to the manufacturer’s instructions. Supernatant (4 μl) was mixed with a color and buffer reagent mixture (250 μl) and the absorbance was read at 570 nm on a plate reader (FMAX, N=4). All ALP activity and calcium deposition measurements were normalized to DNA content. To measure GAG production, chondrogenically differentiated 3D printed hMSC constructs were digested in papain buffer (1 mL, pH 6.5)) containing papain (25 μg m/l, Sigma), l-cysteine (2 × 10^-3^M, Sigma), sodium phosphate (50 × 10^-3^M) and EDTA (2 × 10^-3^M) at 65 °C overnight. GAG content (N=4) was quantified by a dimethylmethylene blue assay ^4^ and DNA content (N=4) was measured using the PicoGreen assay as described above. GAG content was also normalized to DNA content. Quantitative data were expressed as mean ± standard deviation. Statistical analysis was performed with unpaired Student t-test using Graphpad Prism (GraphPad). A value of p<0.05 was considered statically significant.

## Supplementary figures

**Fig. S1.**
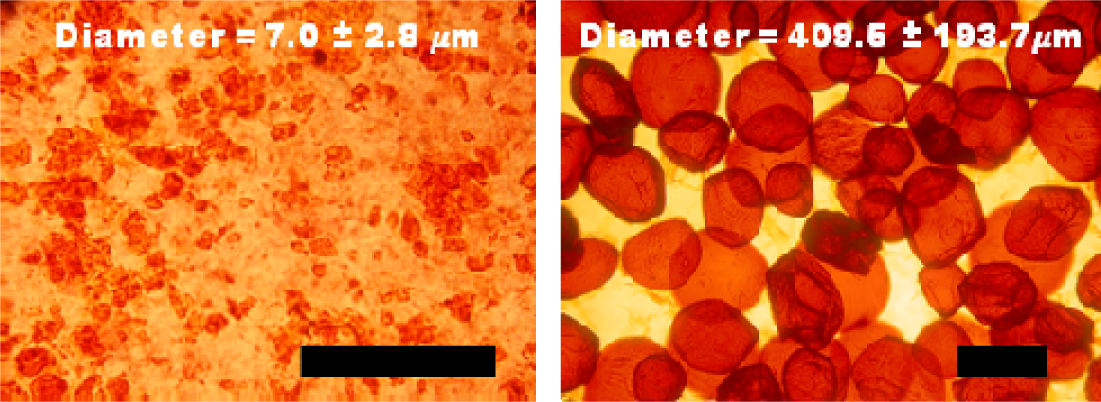
Photomicrographs of Safranin-O stained smaller (left) and larger (right) OMA microgels. Scale bars indicate 200 m.

**Fig. S2.**
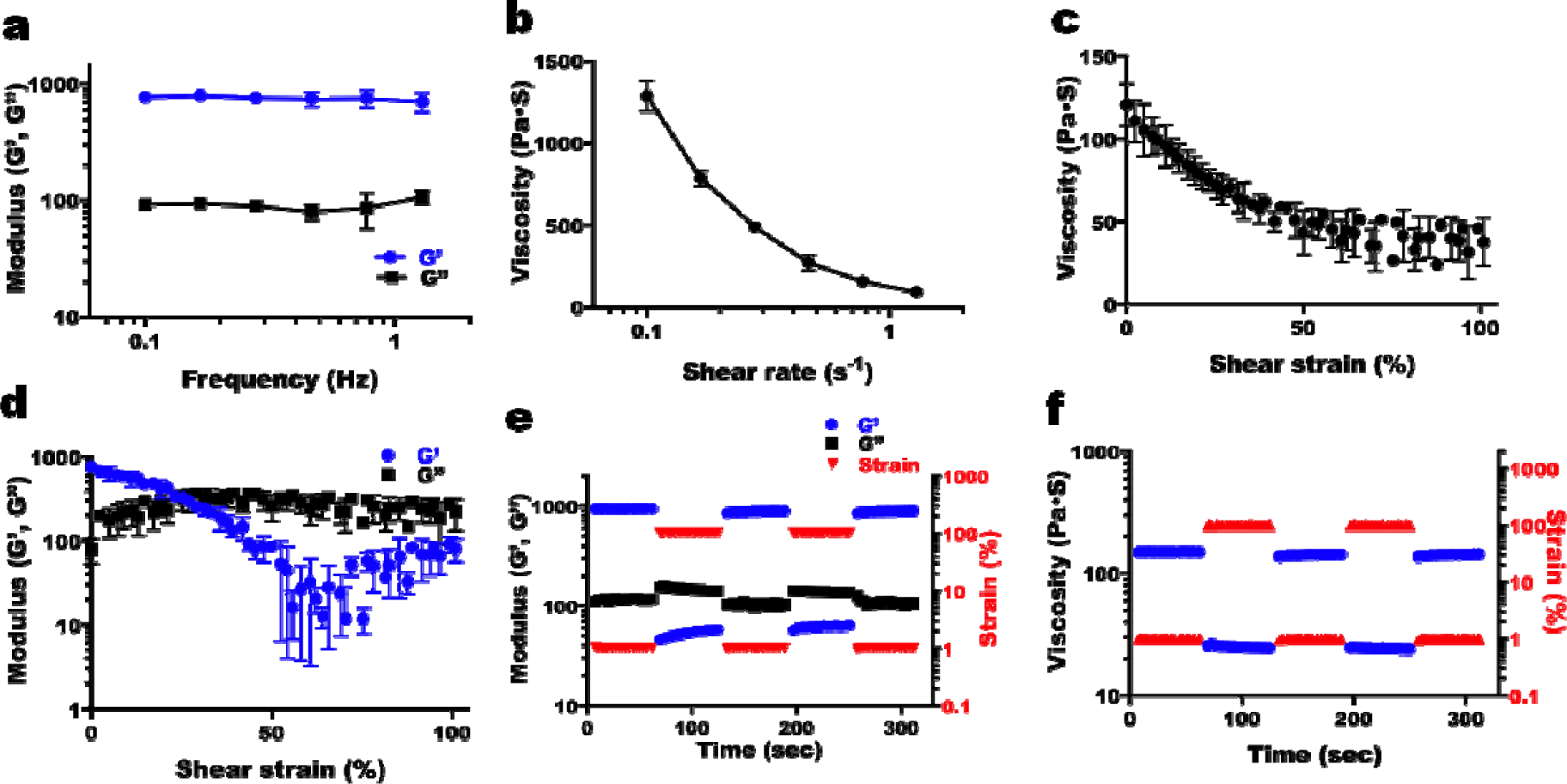
**(a)** Storage (G’) and loss (G”) moduli of alginate microgel supporting medium (mean microgel diameter = 409.6 ± 193.7 μm) as a function of frequency. G’ is larger than G” over the measured frequency range and bot moduli exhibit frequency independence. Viscosity measurements of alginate microgel supporting medium as a function of **(b)** shear rate and **(c)** shear strain demonstrate its shear-thinning behavior. **(d)** G’ and G” of the alginate microgel supporting medium as a function of shear strain exhibit its shear-yielding behavior and gel-to-sol transitio at higher shear strain. **(e)** Shear moduli and **(f)** viscosity changes in dynamic strain tests of the alginate microgel supporting medium with alternating low (1%) and high (100%) strains at 1 Hz demonstrate its rapid recovery of strength and viscosity within seconds, which indicates “self-healing” or thixotropic properties.

**Fig. S3.**
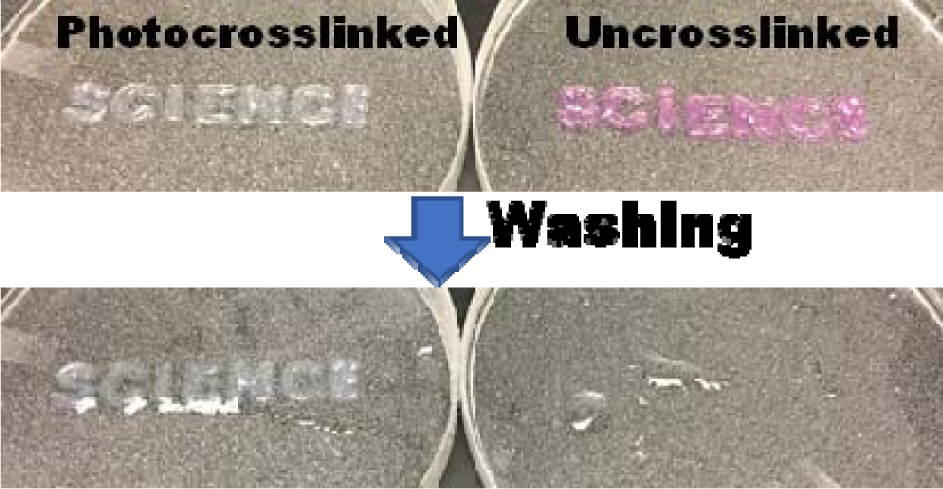
Optical images of photocrosslinked and uncrosslinked microgels before and after washing process.

**Fig. S4.**
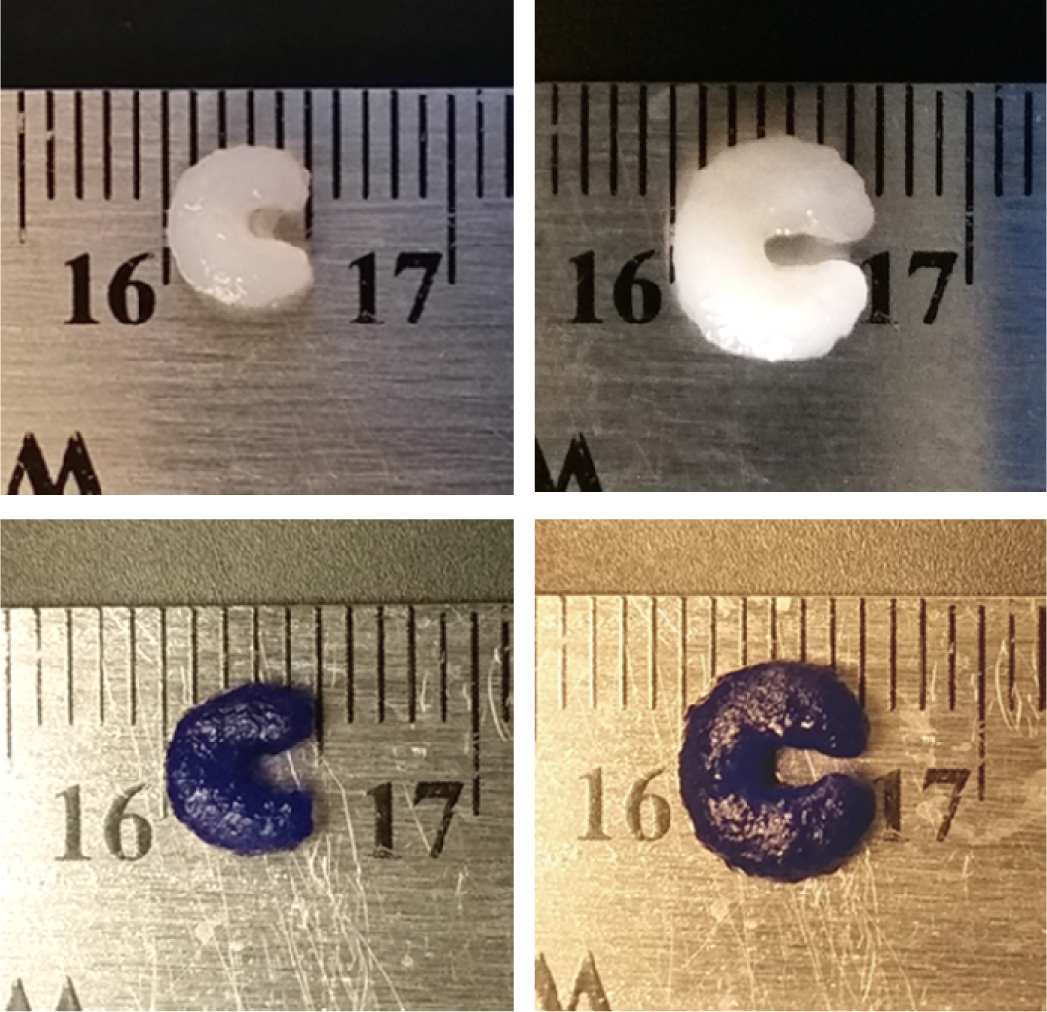
Optical images of chondrogenically differentiated hMSC constructs **(e)** before and **(f)** after Toluidine blue O staining (letter C).

**Fig. S5.**
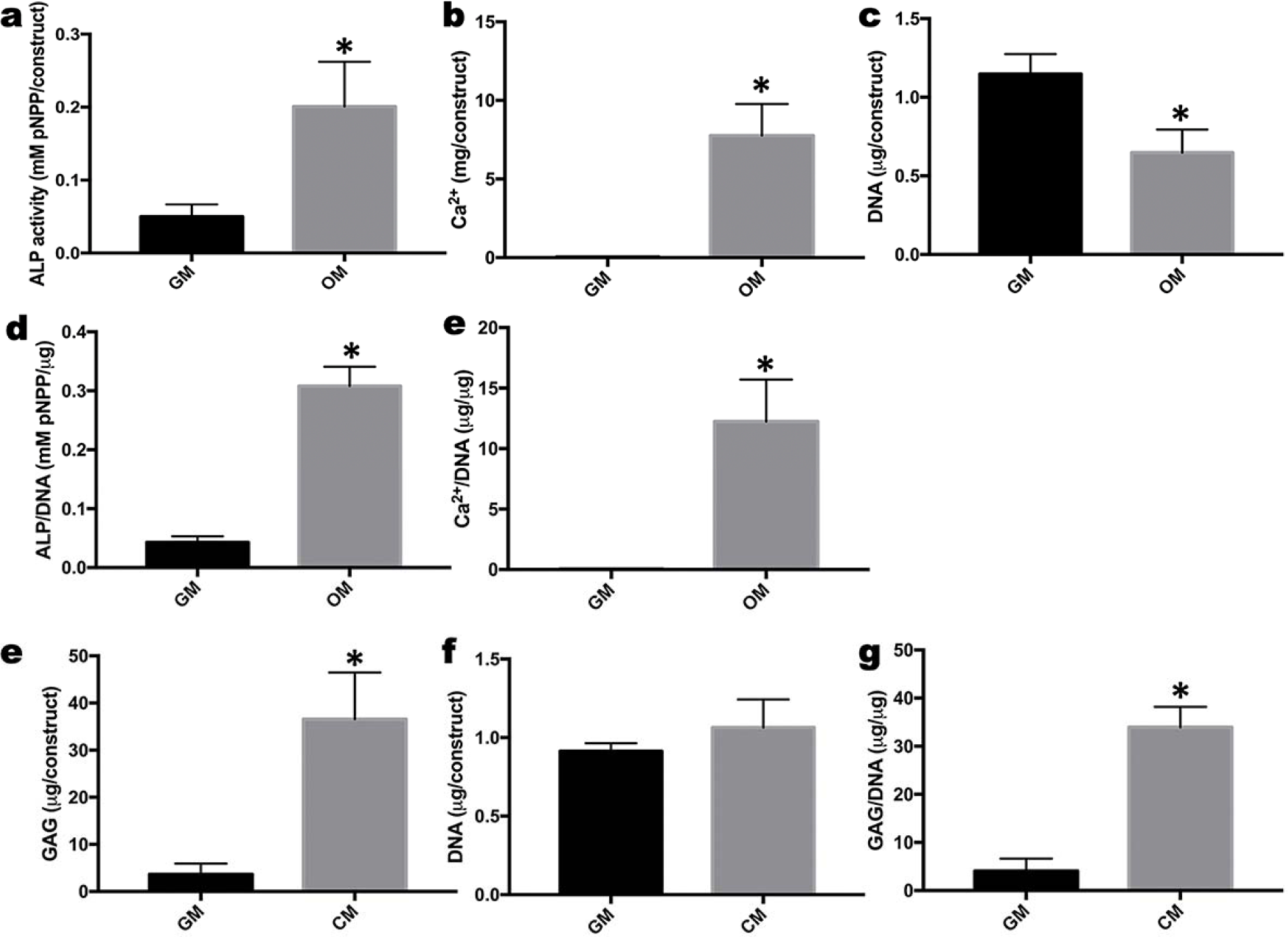
Quantification of (a) ALP activity, (b) Ca^2+^ and (c) DNA content in the 3D printed hMSC constructs cultured in growth media (GM) and osteogenic media (OM) for 4 weeks. (d) ALP activity and (e) Ca^2+^ normalized by DNA. Quantification of (e) GAG production and (f) DNA content in the 3D printed hMSC constructs cultured in growth media (GM) and chondrogenic media (CM) for 3 weeks. (g) GAG content normalized by DNA. *\*p*<0.05 compared to GM.

**Fig. S6.**
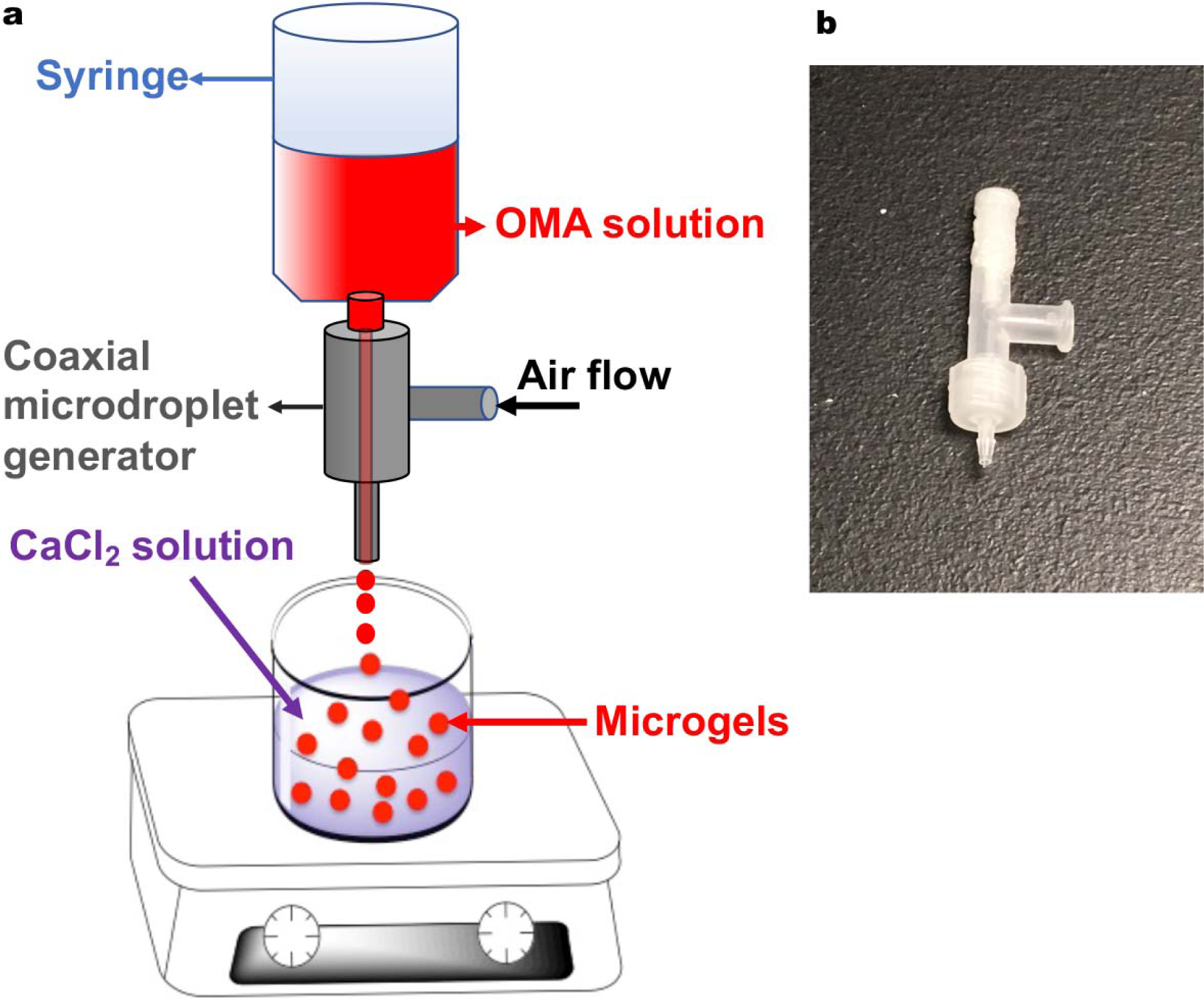
(a) Schematic diagram of coaxial airflow-induced microgel generating system and (b) a photograph of the custom coaxial airflow-induced microdroplet generator.

**Movie S1.** Bioprinting of the letter “C” using a living stem cell-only bioink into an alginate microgel supporting medium.

**Movie S2.** Bioprinting of an ear using a living stem cell-only bioink into an alginate microgel supporting medium.

**Movie S3.** The printing needle can freely move within an alginate microgel supporting medium without creating crevasses.

